# Multifunctional optrode for opsin delivery, optical stimulation, and electrophysiological recordings in freely moving rats

**DOI:** 10.1101/2021.04.30.441836

**Authors:** Kirti Sharma, Zoe Jaeckel, Artur Schneider, Oliver Paul, Ilka Diester, Patrick Ruther

## Abstract

**Objective:** Optogenetics involves delivery of light-sensitive opsins to the target brain region, as well as introduction of optical and electrical devices to manipulate and record neural activity, respectively, from the targeted neural population. Combining these functionalities in a single implantable device is of great importance for a precise investigation of neural networks while minimizing tissue damage.

**Approach:** We report on the development, characterization, and in vivo validation of a multifunctional optrode that combines a silicon-based neural probe with an integrated microfluidic channel, and an optical glass fiber in a compact assembly. The silicon probe comprises an 11-μm-wide fluidic channel and 32 recording electrodes (diameter 30 μm) on a tapered probe shank with a length, thickness, and maximum width of 7.5 mm, 50 μm, and 150 μm, respectively. The size and position of fluidic channels, electrodes, and optical fiber can be precisely tuned according to the in vivo application.

**Main results:** With a total system weight of 0.97 g, our multifunctional optrode is suitable for chronic in vivo experiments requiring simultaneous drug delivery, optical stimulation, and neural recording. We demonstrate the utility of our device in optogenetics by injecting a viral vector carrying a ChR2-construct in the prefrontal cortex and subsequent photostimulation of the transfected neurons while recording neural activity from both the target and adjacent regions in a freely moving rat. Additionally, we demonstrate a pharmacological application of our device by injecting GABA antagonist bicuculline in an anesthetized rat brain and simultaneously recording the electrophysiological response.

**Significance:** Our triple-modality device enables a single-step optogenetic surgery. In comparison to conventional multi-step surgeries, our approach achieves higher spatial specificity while minimizing tissue damage.

**Graphical abstract:** 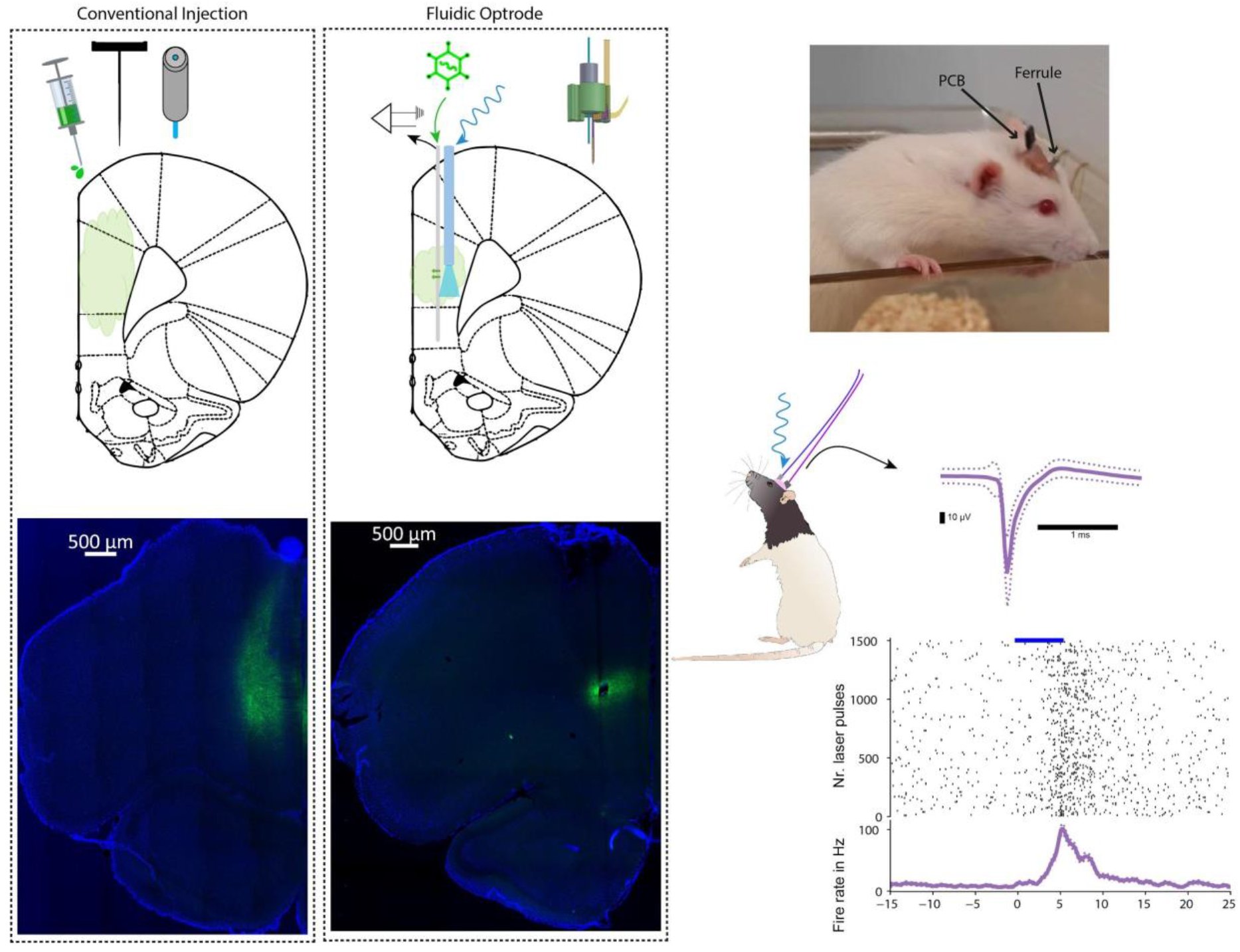

## 1. Introduction

One of the main questions of neuroscientific research is what neural activity underlies a specific behavior or function. While research has advanced with techniques such as the correlation of neurophysiological recordings with behavior, neuromodulation enables a closer observation of causal or necessary neural functions for a specific behavior. Neuromodulation with high spatiotemporal resolution is vital to efficiently investigate neural networks. What was once accomplished with high-artifact electrical stimulation is now often investigated via a more specific, less artifact-prone optical neuromodulation termed optogenetics (Boyden et al., 2005).

This technique is a combination of genetic and optical methods which allows to reversibly excitate or inhibit activity in specific cell populations in systems as complex as freely moving animals (Deisseroth, 2011). This involves integration of light-sensitive membrane proteins, i.e., opsins, into neurons via genetic manipulation. Opsins react to specific wavelengths of light, allowing for a manipulation of neuronal activity. Some opsins are depolarizing, such as channelrhodopsin-2 (ChR2), a cation channel that induces action potentials in response to blue light (Boyden et al., 2005; Nagel et al., 2003). Others are hyperpolarizing, such as the chloride pump halorhodopsin (Zhang et al., 2007). Integration of opsins into neurons has enabled optic neuromodulation at high spatiotemporal resolution (Boyden et al., 2005; Han & Boyden, 2007). This field encompasses diverse strategies to target specific cell populations or projections, making it a powerful toolbox for dissecting neural circuits (De La Crompe et al., 2020). The genetic introduction of opsins is achieved via viral vectors with cell-type-specific promoters, transgenic animals, or a combination of both. This provides specific targeting of defined cell-types or projection-specific neural populations. The desired modulation target can then be further spatially restricted by the area of light application. The type of opsin integrated and light pattern applied can also be modified to provide high temporal control over the neuromodulation (Li et al., 2005). In combination, these adaptable features offer the benefit of high spatiotemporal neural inhibition or excitation of specific cell populations or projections, proving advantageous over electrical stimulation, which can only unspecifically activate all cells in the area near the stimulation site with comparably low spatiotemporal resolution.

While optogenetics currently is the prominent method of neuromodulation, electrophysiology is still the method of choice for recording neural activity with the highest temporal resolution. As opposed to electrical stimulation, optogenetic modulation causes little interference with electrophysiological recordings. Neuroengineering has made significant progress in applying these two methods for simultaneous neuromodulation and recording, for example, via optrodes (H. Kim et al., 2020; K. Kim et al., 2020; Royer et al., 2010). Optrodes, which combine the functions of light application and electrophysiological recording, have become increasingly relevant for optogenetic experiments in order to optogenetically alter a specific cell population while recording its neuronal activity.

Despite its undisputed merits, the technique of optogenetics leaves room for further refinements. For example, one way of integrating opsins into an in vivo model relies on viral injection into a brain area. This is typically achieved with stereotaxic surgery using a nanosyringe or micropipette. After viral injection, the syringe is retreated from the brain, whereupon another device must be inserted in order to provide a light source for accomplishing the desired optogenetic manipulation. This can pose problems, such as backflow of the virus when retreating the injection tool (Fig. S1), or damage to the brain tissue due to repeated device insertions in the same area. To avoid these issues and allow for a single-insertion implantation surgery, it is of interest to combine fluidic, optical and electrical recording functionalities in a single device. Recently such triple-modality probes have come into focus for more efficient interrogation of neural circuitry (Canales et al., 2015; Park et al., 2017; Shin et al., 2019, 2021). Park et al. 2017 and Canales et al. 2015 integrated these modalities in flexible polymer fibers fabricated using a fiber drawing process. They made large strides with long-term, complex experimental testing of their devices in mice with simultaneous optical stimulation, neural recording, and viral (Park et al. 2017) or drug delivery (Canales et al. 2015). Although these multifunctional fiber probes come with a miniature footprint, the technology suffers from a few design limitations. The position of electrodes, fluidic outlets and optical ports is limited to the tip of the fibers, and a manual back-end connectorization of individual electrodes limits the expansion of the number of recording sites. Microelectromechanical systems (MEMS) technology offers higher design flexibility while maintaining a small footprint, as shown in silicon-based multifunctional probes by Shin et al. 2019. They monolithically integrated an optical waveguide, fluidic channel, and electrodes in a planar multi-shank array. Shin et al. 2019 demonstrated the functionality of their probe through simultaneous drug delivery, optical stimulation and neural recording in transgenic mice through acute experiments, but did not implement the probe for viral delivery or electrophysiology in freely moving animals. In this work, we present a multifunctional optrode which combines a MEMS-fabricated microfluidic probe with an optical fiber, and implement it for viral delivery, and subsequent optical stimulation and neural recording in a chronically implanted freely moving rat.

MEMS-based neural probes can be equipped with microfluidic channels using three different fabrication approaches, i.e., surface micromachining, substrate bonding, and bulk micromachining (Sim et al., 2017). Surface micromachining results in channels on top of the device substrate. Typically, the fabrication process starts with patterning a sacrificial material which is subsequently covered by a channel material such as silicon nitride (Retterer et al., 2004) or parylene C (Neeves et al., 2006), followed by the selective removal of the former sacrificial material to realize microchannels. Particularly in the case of long and narrow channels, the channel formation is restricted by the diffusion-limited removal of the sacrificial material from the channel inlet and outlet ports. As a solution, channels can be equipped with etch holes along their length that facilitate the diffusion-driven etching of the sacrificial material (Retterer et al., 2004). However, these etch holes need to be subsequently sealed using a vapor phase deposition process. In the case of the wafer bonding approach, recessed channels are created in one substrate wafer; they are subsequently sealed from the top side by bonding a second material to the substrate. Silicon (Seidl et al., 2010; Spieth et al., 2011), glass (Shin et al., 2019), parylene (Ziegler et al., 2006) and polyimide (Moser et al., 2012) have been applied for channel sealing. Up to this point, the channel dimensions are not constrained by any processing step. However, following the critical wafer bonding, an additional thinning process might be required (Seidl et al., 2010; Shin et al., 2019; Spieth et al., 2011). Lastly, bulk micromachining is capable of generating buried microfluidic channels in silicon (Si) wafers by applying Si etching and subsequent sealing processes (Groenesteijn et al., 2017). Channels are formed by either wet (J. Chen et al., 1997; Cheung et al., 2003) or dry (P.-J. Chen et al., 2006; Dijkstra et al., 2007; Pongrácz et al., 2013) etching of Si through small openings in a micro-patterned masking layer. The channel size can be controlled by adjusting the etching time. The small openings in the masking layer are subsequently sealed using chemical vapor deposition (CVD) processes. In this work, we used bulk micromachining of Si using continuous-flow (cf) xenon difluoride (XeF_2_) etching to produce buried channels with semi-circular cross-sections in the Si substrate. XeF_2_ etching of Si is an isotropic, gas-phase, plasma-free process performed at room-temperature; it offers a very high etch selectivity to materials commonly used in MEMS engineering such as silicon oxide (SiOx), silicon nitride (SiNy), photoresist (PR), and aluminum (Williams et al., 2003). We have characterized cf-XeF_2_ etching to realize microfluidic channels in detail in our previous work (Sharma et al., 2019).

In this work, we developed a microfluidic neural probe with a buried fluidic microchannel and a high-density linear array of recording electrodes at application-specific locations along the probe length. Furthermore, we created the multifunctional optrode by assembling the microfluidic probe close to an optical fiber in a small, 3D-printed unit. The optical fiber interfaced with an external laser offers a wide range of stimulation wavelengths and optical powers. The microfluidic probe and the fiber tip are aligned in such a way that specific fluidic injection and light illumination are allowed in either the prelimbic cortex (PL) or infralimbic cortex (IL), while it is possible to record from both areas. These areas are adjacent yet separate subareas of the medial prefrontal cortex, which are also heavily interconnected (Mukherjee & Caroni, 2018). The reciprocal connectivity between PL and IL has been shown to affect IL-mediated learning (Mukherjee & Caroni, 2018). It has further been shown that PL and IL play opposing roles in various behaviors such as goal-directed learning, fear learning, and learning vs applying rules (Fenton et al., 2014; Hardung et al., 2017; Mukherjee & Caroni, 2018). Because of their opposing functions, it is important to specifically target one area without externally influencing the other; only then can clear behavioral results be acquired. This can prove difficult, since injections into small areas such as IL can also end up transducing neighboring areas, such as the PL. This can occur due to general viral spread, but also from backflow of the virus when retreating the injection needle from the brain, as demonstrated in Fig. S1. Considering the interconnectivity, proximity, and context-dependent functionality of these two areas, it is of interest to target one area while recording in both.

With fluid delivery, light delivery, and recording electrodes combined in a single device, our multifunctional optrode enables a single-insertion implantation surgery for optogenetic application with neurophysiological recording. We demonstrate, to our knowledge, the first utility of a device for injecting the viral solution, chronic implantation for several weeks to allow viral expression, and photostimulating neurons near the injection site while recording neural activity both at and adjacent to the injection area in an awake, freely moving rat. Additionally, this novel tool can also be used for delivery of other pharmacological agents in the brain. In another experiment, therefore, we modulated neural activity in an anesthetized rat through bicuculline injections and simultaneously recorded the electrophysiological response.

## 2. Materials and Methods

### 2.1 Device design

The multifunctional optrode comprises a microfluidic neural probe combined with a ferrule-terminated optical glass fiber. The probe and the glass fiber are aligned in parallel and close to each other on a 3D-printed unit (Fig. 1(A)). A schematic of the fluidic probe is depicted in Fig. 1(B). It contains a fluidic channel and recording electrodes along its slender probe shank. The shank is attached to a broader probe base that mediates the fluidic and electrical interconnections via inlet ports and bonding pads, respectively. The protruding design of the fluidic inlet port facilitates the in-plane connection of flexible tubings. The electrodes are connected to the external recording equipment using a highly flexible polyimide (PI) ribbon cable flip-chip bonded to the contact pads (Kisban et al., 2009). The cross-sections of the probe shank along lines A-A’ and B-B’, as illustrated in Fig. 1(B), are shown in Fig. 1(C(viii)). As indicated, the fluidic channel is located beneath the probe surface directly under the electrodes and interconnection lines. The fluidic outlet of the channel opens to one side of the probe shank.

**Figure 1:**
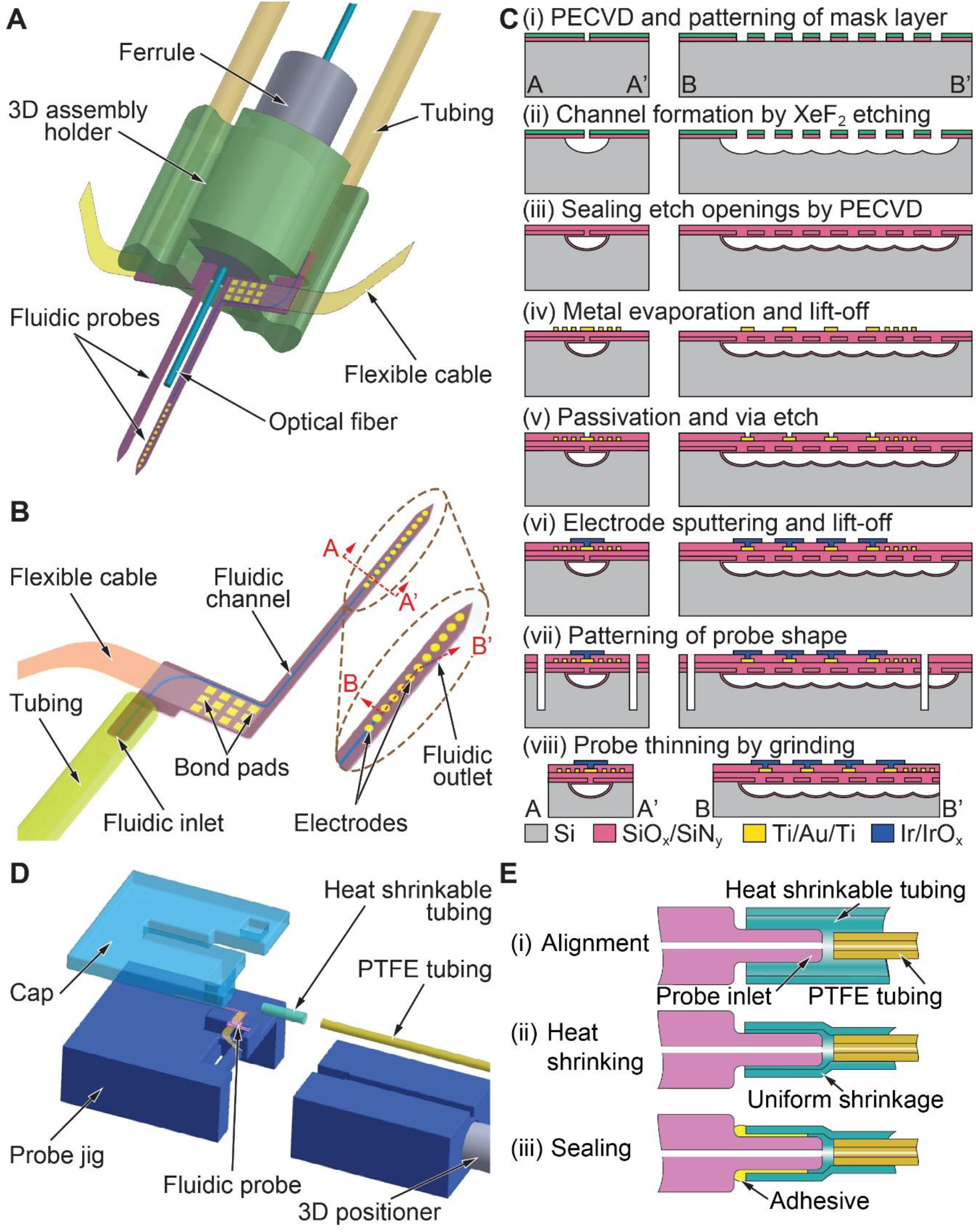
(A) Schematic of multifunctional optrode based on a microfluidic neural probe and an optical glass fiber assembled on a 3D-printed unit. If needed, multiple probes can be arranged around a single optical fiber. (B) Schematic of the microfluidic neural probe illustrating the probe shank carrying a fluidic channel and recording electrodes, and the probe base interfacing a flexible fluidic tubing and PI-based ribbon cable. (C) Probe fabrication process based on cf-XeF_2_ etching of bulk Si. It illustrates the probe shank cross-sections along lines A-A’ and B-B’ in (B). (D) Assembly jig used to attach the PTFE tubing to the fluidic probe and (E) assembly sequence of attaching fluidic tubing to fluidic inlet of Si-based probe.

For the application chosen in this study, we designed Si-based fluidic probes comprising a single 11-μm-wide fluidic channel and 32 recording electrodes (diameter 30 μm) on a tapered probe shank with a length, thickness, and maximum width of 7.5 mm, 50 μm, and 150 μm, respectively. The electrodes are arranged at a pitch of 60 μm towards the tip (electrodes 1-16) and a pitch of 85 μm towards the upper part of the shank (electrodes 17-32). This was custom designed so that we have 16 recording electrodes in each area, i.e., the IL and the PL. Each probe carries a single fluidic channel that ends in two closely spaced outlet ports. There are two design variants of the fluidic probe, which vary in the position of the fluidic outlet. The IL configuration has fluidic outlet ports in the lower part of the shank (at 345 μm and 445 μm from the tip), while the PL configuration has fluidic ports in the upper part of the shank (at 1845 μm and 2045 μm from the tip). We used an optical glass fiber that is 225 μm in diameter and has a numerical aperture of 0.37 at a core diameter of 200 μm. The tip of the fiber is positioned roughly 100 μm above the fluidic ports. The combination of a fiber with either of the two fluidic probe variants allows us to inject and optogenetically target one area (PL or IL) while recording neural activity from both areas.

### 2.2 Multifunctional optrode fabrication and assembly

#### a. Microfluidic probe

The fabrication process of the microfluidic probes is schematically summarized in Fig. 1(C). It is carried out on 4-inch, 525-μm-thick, single-side polished Si wafers which comprise dry etched alignment marks on the wafer frontside. These marks ensure the precise alignment of multiple photolithography masks used in further processing steps, as detailed below.

##### Fluidic channel patterning

The wafer frontside and the alignment marks are first covered by a 460-nm-thin stack of high-frequency (HF) SiO_x_/SiN_y_/SiO_x_ layers with low tensile mechanical stress realized using a plasma-enhanced chemical vapor deposition (PECVD) process. This layer stack serves as a masking layer during the subsequent etching of Si using XeF_2_. A linear array of small etch openings (footprint 2×4 μm^2^) arranged at a pitch of 10 μm is transferred into the SiO_x_/SiN_y_/SiO_x_ masking layer using photolithographically structured photoresist (PR) (AZ1518, MicroChemicals GmbH, Ulm, Germany) and reactive ion etching (RIE), as shown in Fig. 1(C)(i). The PR is not stripped after the RIE sequence and acts as an additional masking layer during the subsequent cf-XeF_2_ etching. Before XeF_2_ etching, the wafers are dipped in 1% hydrofluoric acid (HF) for 10 s and dried on a hotplate at 120 °C for 60 s. This wet etching is essential for removing any RIE process-related residues or native silicon oxide from the etch openings, providing a clean Si surface. The wafers are then directly transferred into the cf-XeF_2_ etch system from Memsstar Ltd. (Livingston, UK). This system operates in a continuous-flow configuration, where the nitrogen (N_2_) carrier gas flows continuously through a container comprising XeF_2_ crystals, thus propelling XeF_2_ vapor into the etch chamber. The etch chamber maintains a constant pressure of 2.7 mbar during the entire etching process (Drysdale et al., 2015).

The XeF_2_ vapor isotropically etches Si through the small etch openings of the masking layer. Once the undercut length exceeds 3 μm, the adjacent etch fronts merge to form a long channel with a semi-circular cross-section along A-A’ while a slightly corrugated channel bottom results along B-B’ (Fig. 1(C)(ii)). Etching the wafers for 60 s results in fluidic channels with a width of 15 μm. The perforated nature of the thin channel cover prohibits the use of ultrasound-based processes as they would damage these thin dielectric membranes. As a consequence, the PR is next stripped using a plasma-based process.

The cf-XeF_2_ etching of bulk Si is followed by the deposition of an 8-μm-thick, stress-compensated HF PECVD SiO_x_/SiN_y_ layer stack to seal the etch openings (Fig. 2(C)(iii)). This results in the channel sidewalls being covered with the same materials, which terminates once the etch openings are sealed. As a result, the final width of the fluidic channel is reduced from 15 μm to about 11 μm. Despite the fact that 4-μm-thin layer stacks were found to be sufficiently thick to seal the 2×4 μm^2^ large etch openings, thicker stacks are applied as they result in a smoother top surface. This is a requirement for reliably patterning the thin metal lines, which interface the electrodes with the contact pads, at minimal width and space values across the sealed channels. A thicker stack also compensates for variations in the deposition rate across the wafer that prevent a precise control of the final thickness of the deposited layer stack.

**Figure 2:**
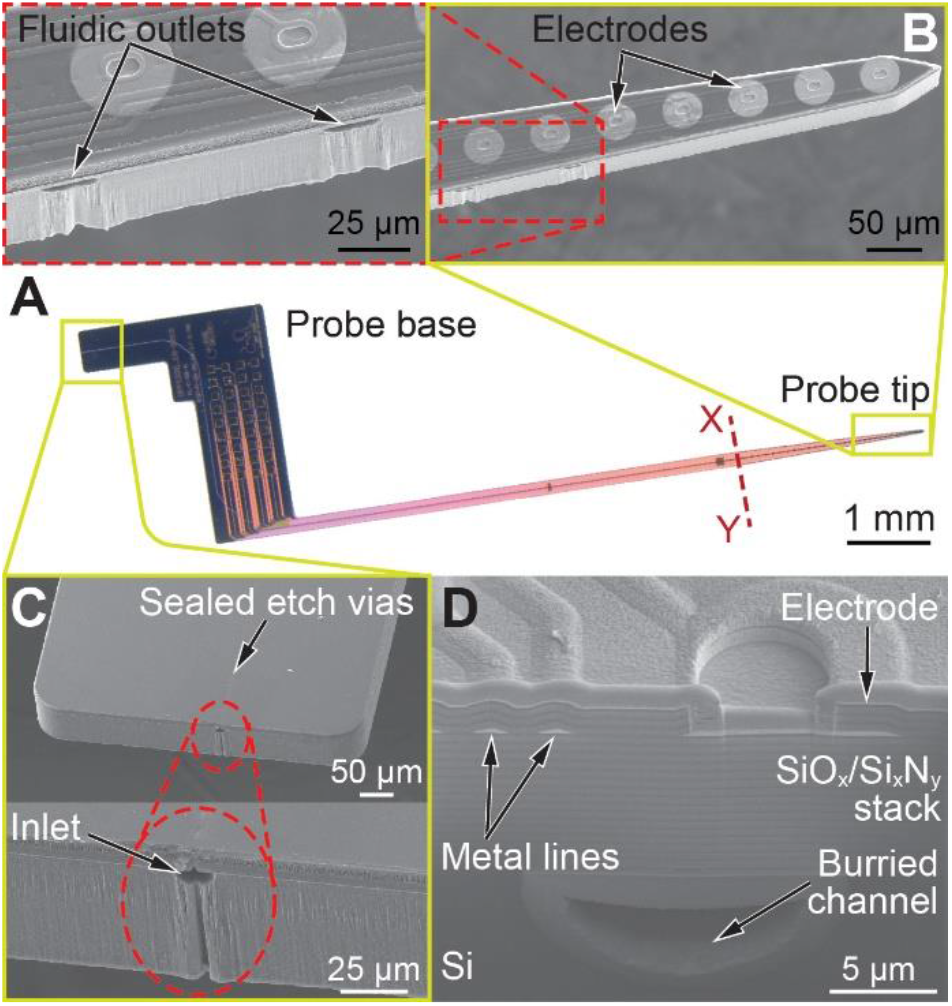
Fabricated 50-μm-thick microfluidic probe. (A) Fluidic probe with base and a 7.5-mm-long and maximally 150-μm-wide tapered shank. (B) Scanning electron microscopy (SEM) micrographs of probe tip with recording electrodes and fluidic outlets. (C) SEM micrographs of fluidic inlet at the probe base. (D) Cross-section of fluidic probe along X-Y given in (A) realized using focused ion beam (FIB) etching showing the buried fluidic channel in Si as well as metal interconnection lines and electrode over the surface sealed by layer stacks of SiO_x_/SiN_y_

At this stage, fluidic channels buried in the Si wafer under a planar dielectric layer have been realized. Further steps for electrode patterning and probe shaping are based on the process described elsewhere in detail (Herwik et al., 2009).

##### Metallization and electrode patterning

Next, bond pads and interconnection lines (line/space 1.5 μm/1.5 μm) as well as disks at the position of the electrodes are defined by a metal layer stack of titanium (Ti)/gold (Au)/Ti (thicknesses 30/250/30 nm). This layer stack is deposited and patterned through evaporation and a lift-off process which is based on the image reversal PR AZ5214E (Microchemicals GmbH, Ulm, Germany) (Fig. 1(C)(iv)). The Ti layers act as adhesion promoters for the dielectric layers, while Au is applied to minimize the electrical impedance of the interconnection lines. The metal stack is then passivated by depositing another 1.5-μm-thick, stress-compensated HF PECVD SiO_x_/SiN_y_ layer stack. This passivation is opened at the positions of the electrodes and bond pads using RIE. Next, the underlying Au layer is exposed by etching the top Ti layer in 1% HF, as illustrated in Fig 1(C)(v).

Thereafter, the recording electrodes with a diameter of 30 μm are defined by applying a two-layer lift-off process and sputter-depositing a layer stack of Ti/platinum (Pt)/iridium (Ir)/iridium oxide (IrO_x_) with thicknesses of 30, 150, 100 and 200 nm, respectively. The lift-off process applies a combination of the PRs LOR5A (Kayaku Advanced Materials, Westborough, MA, USA) and AZ1518 (Microchemicals GmbH, Ulm, Germany) (Fig. 1(C)(vi)). The Ti and Ir layers serve as adhesion promoters to the underlying passivation and Pt, respectively. The Pt layer acts as the electrode base material, while the IrO_x_ coating is implemented to lower the electrode impedance.

##### Probe patterning

Finally, the etching before grinding (EBG) process (Herwik et al., 2011) is used to realize thin fluidic probe shanks and release them from the Si substrate. First, a combination of photolithography, RIE, and deep reactive ion etching (DRIE) is used to define the in-plane geometry of the microfluidic probes (Fig. 1(C)(vii)). In this process sequence, RIE opens the about 10-μm-thick stack of dielectric layers, while DRIE creates 60-μm-deep trenches inside the Si wafer. These trenches also provide access to the fluidic channels by defining in-plane inlets and outlets at the probe base and along the probe shaft, respectively. Finally, wafers are ground from the wafer rear to the desired probe thickness of 50 μm utilizing the grinding service of DISCO HI-TEC Europe GmbH (Kirchheim, Germany) (Fig. 1(C)(viii)).

Due to the fact that the fluidic inlets and outlets are exposed during the grinding process, any residue entering the channels at these positions potentially results in blocked probe channels. On testing fluidic probes after their assembly, we found that almost 50% of the probes had blocked channels. We resolved this issue by soaking the unassembled probes in a dimethyl sulfoxide (DMSO) based organic solvent (TechniStrip Micro D350, Microchemicals GmbH, Ulm, Germany), for an extended period of time during which the solvent was occasionally heated to 70 °C. As a consequence of the additional cleaning, more than 90% of the probes were fluidically functional. Alternatively, we spin-coated PR to the DRIE processed wafer frontside prior to grinding. In this way, the channel inlets and outlets are covered by PR to prevent their contamination with fine particles of Si residue. After grinding, probes were soaked as well in TechniStrip Micro D350 for at least one week to ensure a complete removal of PR from the channels. This technique also resulted in over 90% fluidically functional probes. It is important to note that there are additional reasons for channel blockages observed during testing, such as dust contamination during probe assembly or testing.

#### b. Optical fiber

Multimode optical fibers (225 μm diameter, 0.37 NA, Thorlabs, Newton, NJ, USA) were cut with a diamond knife and glued into stainless steel ferrules (2.5 mm outer diameter, 230 μm bore diameter, Thorlabs) with heat-solidifying epoxy (Precision Fiber Products, Chula Vista, CA, USA). They were polished on one side with 5 μm grit silicon carbide paper, 3 μm grit aluminum oxide paper and finally 0.3 μm grit calcined alumina paper (Thorlabs). The fibers were cut down with a diamond knife to 6 mm for the PL-configuration and 7.2 mm for the IL-configuration probe, so that the tip of the fiber was ca. 100 μm above the fluidic ports when assembled next to the microfluidic probe. They were also tested for light transmission by measuring their optical output power with a power meter (Model PM100D, Thorlabs).

#### c. Multifunctional optrode assembly

The assembly process of the multifunctional optrode establishes on the one hand the electrical and fluidic interfaces to the silicon microfluidic probe, and on the other hand the alignment of the assembled probe relative to the ferrule-terminated optical fiber on the 3D-printed unit, as schematically shown in Fig. 1(A).

##### Microfluidic probe assembly

First, the microfluidic probe is bonded to a custom-made, highly flexible PI ribbon cable employing a flip-chip-bonder (Fineplacer 96λ, Finetech, Berlin, Germany). The ribbon cables with Au-electroplated contact pads are realized, as described in detail elsewhere (Kisban et al., 2009). By applying force, temperature, and ultrasonic power, Au-based contact pads on both the probe base and the cable are joined. The probe-cable interface around the contact pads is underfilled with a two-component epoxy (Epotek 301-2, Epoxy Technology, Billerica, MA, USA) cured at 65°C for 2 hours to encapsulate the contact pads against any liquid to which the probe is exposed during the in vivo experiments. Next, a polytetrafluorethylene (PTFE) tubing (inner diameter (ID) 300 μm, outer diameter (OD) 760 μm) is bonded to the in-plane inlet of the probe, enabling fluidic interfacing to the external fluidic equipment, e.g. a syringe pump, for fluidic supply. As initially proposed by Spieth et al. 2009, a short piece of heat-shrinkable (HS) tubing (Raychem MT 2000, Schaffhausen, Switzerland) is used to establish this connection at minimal dead-volume (Spieth et al., 2009). The assembly procedure is carried out using custom-made tools, as schematically illustrated in Fig. 1(D). The protruding part of the probe base and PTFE tubing, surrounded by the HS tubing, are aligned in a straight line (Fig. 1(E)(ii)). By applying the air stream of a hot air gun at 180 °C, the HS tubing is uniformly shrunk around both the probe inlet and the PTFE tube simultaneously (Fig. 1(E)(ii)). Another epoxy (Epotek 353 ND-T, Epoxy Technology) is applied to fill the remaining gaps around the probe inlet (Fig. 1(E)(iii)). It is cured in an oven at 150°C for 1 hour to achieve a leak-proof connection.

##### Consolidation of fluidic, optical, and electrical units

A 3D-printed cylindrical unit with an external diameter and height of 6 mm and 5 mm, respectively, is used for aligning and fixing the microfluidic probe and the ferrule-terminated fiber close to each other, as schematically shown in Fig. 1(A). It consists of a 4-mm-deep hollow cavity in the centre to hold the fiber ferrule. During assembly, the microfluidic probe is first aligned and glued near the base of the 3D-printed unit using a UV-curable adhesive (U306, Cyberbond Europe GmbH, Wunstorf, Germany). The ferrule-terminated optical fiber is then inserted into the 3D-printed component. By design of this 3D-printed unit the distance between the frontside of the microfluidic probe and the optical fiber is controlled to a targeted lateral separation of around 200 μm. The angular misalignment between the fluidic probe shank and optical fiber was determined to be within 2° and below. The ferrule is secured on both ends of the 3D-printed unit using a self-adhesive resin cement (RelyX Unicem 2, 3M Deutschland, Neuss, Germany). The PI-based ribbon cable is interfaced to a zero insertion force (ZIF) connector on a custom-made printed circuit board (PCB) compatible with the applied recording system. The interface is encapsulated with the epoxy Epotek 353 ND-T. The total weight of the implant including the PCB is 0.97 g. Custom holders were designed to assist in this assembly process and storage of the assembled multifunctional optrodes, as illustrated in Fig. S2.

### 2.3 Electrical and fluidic probe characterization

The electrical impedance of microelectrodes was measured in Ringer’s solution in a two-electrode setup with a Pt counter electrode in the frequency range from 100 Hz to 40 kHz.

The assembled fluidic probes were in addition hydrodynamically characterized by connecting them to a syringe pump (NE-300-ES, New Era Pump Systems Inc., Farmingdale, NY, USA) using commercially available fluidic connectors. The flow rate was increased from 200 to 1000 nl/min in multiple steps, and the fluidic pressure was recorded using a microfluidic pressure sensor (MPS4, Elveflow, France) equipped in the flow path.

In order to investigate the spatial distribution of a liquid injected from these probes as a function of flow rate, we applied 0.6% agarose gel as a suitable brain model (Z.-J. Chen et al., 2004). The microfluidic probe was connected to a 10-μl glass syringe via thin tubings and a union, similar to the fluidic line in the experimental setup in Fig. 3(B). The probe shank was slowly inserted into a block of agarose gel, and 1 μl of 0.22-μm-filtered, 0.1% bromophenol blue solution was injected using the syringe pump NE-300-ES (New Era Pump Systems Inc.). Multiple injections were performed at flow rates of 50, 100, 200, and 400 nl/min, and the spread of dye was monitored with a microscope camera (MC190 HD, Leica Microsystems GmbH, Wetzlar, Germany).

**Figure 3:**
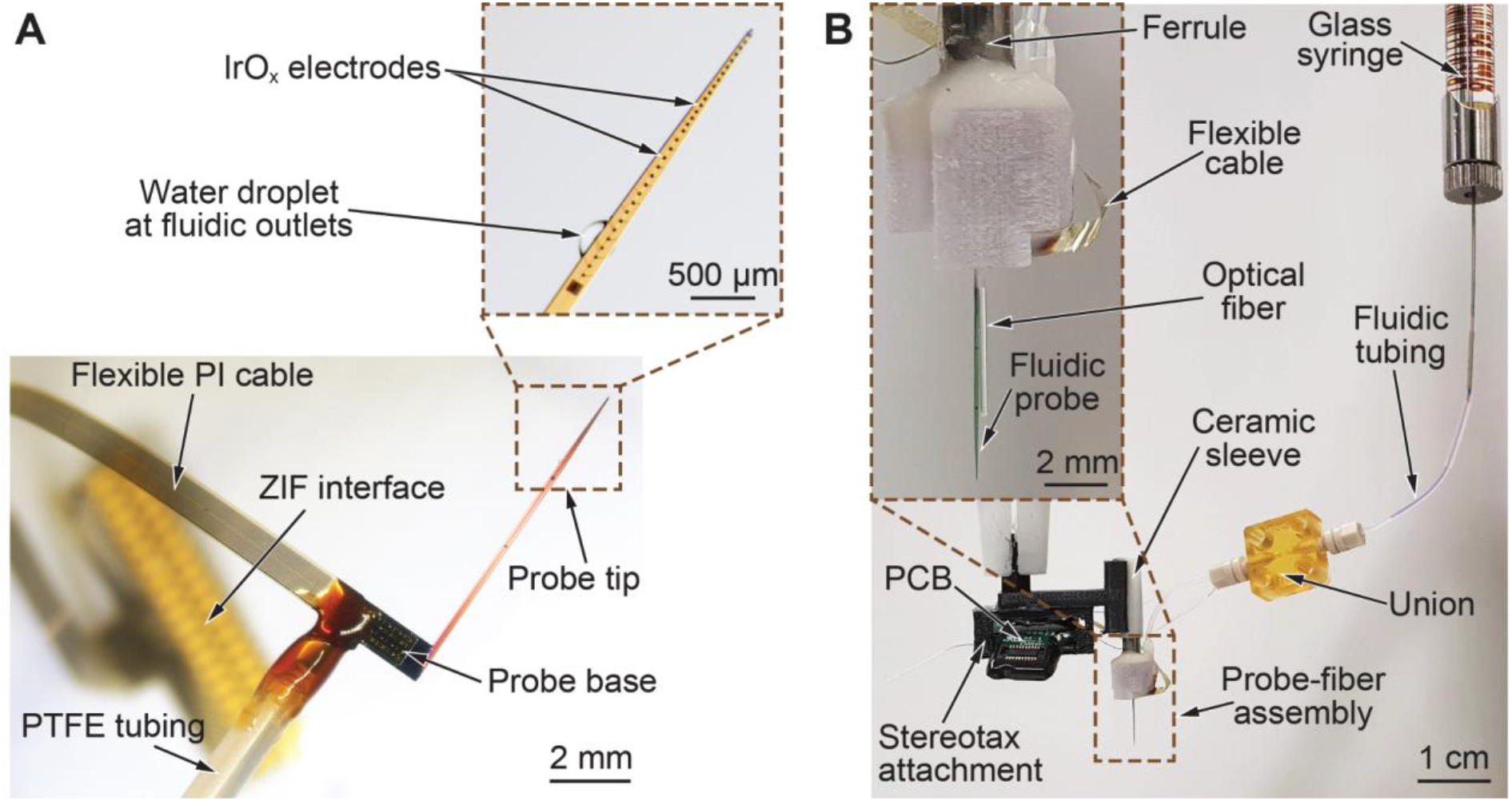
Fabricated fluidic probe and multifunctional optrode: (A) Si-based neural probe assembled to a PTFE tubing and highly flexible PI-based ribbon cable. The fluidic functionality is validated by injecting a small amount of deionized (DI) water through the PTFE tubing. (B) Multifunctional optrode attached to a custom-designed stereotaxic holder. The union is applied as the fluidic interconnection with minimal dead volume between the PTFE tubing of the probe and the tubing interfacing the glass syringe.

### 2.4 In vivo experiment

#### a. Nanosyringe liquid loading

A 10 μl nanosyringe (World Precision Instruments, Sarasota, FL, USA) is first filled with the fluid to be injected before connecting it to the fluidic line of the multifunctional optrode. Since the combined volume of the syringe and the interconnection tubing is much larger than the required injection volume, the syringe is first filled with a saline solution. For this, a 28 gauge microfil (World Precision Instruments) was used to load 7 μl of colored saline into the nanosyringe. A 40-mm-long PTFE tubing (ID 300 μm, OD 760 μm) was then attached to a 26 gauge nanosyringe needle (World Precision Instruments) using a heat shrinkable tubing. The tube-fixed needle was then inserted into the nanosyringe which was then loaded into the stereotaxic holder of the microsyringe pump (World Precision Instruments). The saline solution was pushed to the tip of the PTFE tubing. Next, the subsequent solutions were withdrawn in the following order at 200 nl/min: 1 microliter of corn oil (to prevent the injection fluid from mixing with the saline), 800 nl of the desired injection fluid (in this study bicuculline or ChR2 virus), and 200 nl of air. The open end of the tubing was closed with a silicone cap, and the microsyringe was stored at 4°C until surgery on the following day.

#### b. Nanosyringe connection to multifunctional optrode

The tubings from the nanosyringe and the multifunctional optrode were linked via an interconnect union and fittings (LabSmith, Inc., Livermore, CA, USA). Once connected (Fig. 3(B)), we injected through the nanosyringe in air at a rate of 200 nl/min until a drop of the injection fluid appeared at the outlet of the fluidic probe. We stopped the injection immediately, and removed the drop with a kimtech wipe.

#### c. Stereotaxic surgery for acute recording

All animal related procedures were in accordance with the guideline RL 2010 63 EU and were approved by the Regierungspräsidium Freiburg (approval number TVA G-15-11 and TVA G-20-26). We kept the rats in a reversed 12-hour light cycle, and without any food or water deprivation.

Animals were given an inhalative gas anesthesia with isoflurane using oxygen (O_2_) as a carrier gas. They were then administered intraperitoneal (i.p.) injections of 40 mg/kg ketamine and 0.1 mg/kg medetomidine. Subcutaneous (s.c.) injections of 5 mg/kg carprofen and 0.05 mg/kg buprenorphine were applied prior to surgery. A post-surgery additional injection of carprofen was also applied. Anesthesia was maintained at 0.5 - 2 % isoflurane and 0.5 l/min O_2_. To prevent hypothermia, rats were kept on a heating pad and their temperature was monitored via a rectal sensor. Eyes were kept moist with an ophthalmic ointment (Bepanthen, Bayer Health Care, Leverkusen, Germany). Rats were injected s.c. with 3 ml of saline solution every two hours to maintain their fluid balance. Fur was shaved from the surgical site, and the rats were then head-fixed into a stereotaxic frame (World Precision Instruments,). The surgical surface was disinfected with Braunol (B. Braun Melsungen AG, Melsungen, Germany), followed by Kodan (Schülke, Norderstedt, Germany). A single 2-cm-long incision was made down the skull from anterior to posterior, and the edges of the skin were held apart with surgical clamps. A bone scraper was used to remove the skin tissue from the skull, and 3% hydrogen peroxide was used to thoroughly clean the skull of any leftover debris or tissue. The head was then levelled in the anterior-posterior (AP) direction so that bregma and lambda were within a 0.05 mm height difference of each other. Medial-lateral (ML) levelling was achieved by positioning the head so that ML locations to the right and left of bregma were within 0.05 mm of each other. A craniotomy was drilled around the area of planned injection. One miniature self-tapping screw (J.I. Morris Company, Southbridge, MA, USA) was inserted deep through the skull to serve as a grounding element.

The multifunctional optrode was connected to the stereotaxic frame via a custom designed 3D-printed stereotaxic attachment that is held by a stereotaxic cannula holder (World Precision Instruments) (Fig. 3(B)). A ceramic mating sleeve (2.5 mm diameter, Thorlabs) was glued on one end of the stereotaxic attachment. The ferrule of the multifunctional optrode was inserted into the ceramic sleeve, and the PCB was placed into a slot in the stereotaxic attachment (Fig. 3(B)). A duratomy was made over the injection site, and the ground wire from the PCB was attached to a skull screw. Next, we attached the PCB to a 32-channel data acquisition system using a ZIF-clip headstage (Tucker-Davis Technologies (TDT), Alachua, FL, USA), and inserted the probe to the desired location (AP 3.24 mm / ML 1.0 mm / DV −4.2 mm from surface of brain).

#### d. Acute recordings with bicuculline application

After inserting the probe into the brain, we waited for the tissue to settle around the probe for 5 minutes. We injected 200 nl of 5 mM -(−) bicuculline methiodide (Tocris Bioscience, Bristol, UK) with the syringe pump at 100 nl/min. We recorded before, during, and after the injection to observe the neural effect of the fluidic drug. Recording was performed using a 32-channel data acquisition system (TDT) and filter settings of 300 Hz for high-pass and 5 kHz for low-pass filtering. We acquired neural data with a sampling frequency of F_s_ = 24,414.0625 Hz. After recording, the probe was retreated from the brain and the rat was perfused.

#### e. Chronic implantation

The surgery was similar to the surgery method for the acute recording. Along with one miniature screw for grounding, we placed 4 additional screws into the skull to provide an anchor for the implantation cement. We implanted the probe to the desired depth, and injected 500 nl of pAAV-hSyn-hChR2(H134R)-eYFP-WPRE (University of North Carolina Vector Core, Chapel Hill, NC, USA) at a flow rate of 100 nl/min. Silicone elastomer (Kwik-Cast, WPI, Sarasota, FL, USA) was placed into the craniotomy to prevent brain swelling. We then applied a thin layer of super bond C&B cement (Sun Medical Co., LTD, Moriyama City, Shiga, Japan) between one of the skull screws and around the probe to stabilize the probe in the brain. The PTFE tubing attached to the fluidic probe was cut at the base, and the opening of the tubing was blocked with dental cement. The PCB board was placed at a 40° angle to the base of the skull, and fixed with Paladur (Heraeus, Hanau, Germany). Paladur was further added to create a firm base on the skull. The surgical site was sealed around the implant by interrupted sutures. Rats were fed 5 mg carprofen pellets for three days after surgery, and it was checked during these days that they did not drop to under 80 % of their pre-surgery, ad libitum-food and -water access weight. We waited 6 weeks post-surgery for sufficient viral expression before stimulation experiments.

#### f. Recording and optogenetic stimulation in freely moving animals

The rat was placed in a behavioral box for freely moving exploratory behavior. The box was made from 8-mm-thick acrylic glass and had a size of 45×36×55 cm^3^. For neuronal recordings, broadband signals were simultaneously recorded via ZD32 digital head stages connected via an electrical commutator (ACO64, TDT) to the recording controller (Intan Technologies LLC, Los Angeles, CA, USA). Neural data was recorded at a sampling frequency of 30 kHz. For optogenetic stimulations we connected the rat via a mono mode fiber-optic patch cord to a light source (LightHUB compact laser combiner, OMICRON Laserage, Rodgau-Dudenhofen, Germany) to stimulate with 473 nm pulsed light (5-ms-wide pulses at 10-40 Hz, 1-s-long bursts, 318 mW/mm^2^) while recording neural activity from the transfected and the adjacent non-transfected area.

### 2.5 Data analysis and histology

#### a. Analysis of electrophysiological recordings

We performed the analysis using custom written Python scripts. The broadband signal was high-pass filtered (4. degree butterworth digital, 200 Hz cutoff using scipy.signal.sosfiltfilt). The resulting signal was median subtracted across channels (using 4 channel rolling median window). We defined spikes as time points exceeding 4*std of the median subtracted signal. Spike times were aligned relative to the laser bursts and increase in the firing rate was calculated relative to 1 s period before the laser stimulation. Firing rate was obtained via convolution of the spike times with a Gaussian window. We assessed significance of the modulation of the firing rate via the Bonferroni adjusted ranksum test with prestimulus vs peri stimulus firing rate. For the analysis of neural recordings with bicuculline injection spikes were detected in the same manner directly on high-pass filtered signal; here LFP and spike times were aligned to the start of injection.

#### b. Perfusion and histology

Animals were sacrificed via an i.p. injection of 400 mg/kg sodium pentobarbital (Release®500, WDT, Garbsen, Germany) and a transcardial perfusion of 4 °C PBS, followed by ice-cold 4% paraformaldehyde. The brains were removed and post-fixed for 1-2 days and then transferred to 30% sucrose (Merck KGaA, Darmstadt, Germany) dissolved in PBS, and stored at 4 °C. After the brains were fully saturated with sucrose, we sectioned them into 50-μm-thin slices on a sliding microtome (Leica model SM2010 R, Leica Biosystems Nussloch GmbH, Nussloch, Germany). Images were obtained on an Axioplan 2 Imaging microscope.

## 3. Results

### 3.1 Microfluidic probe fabrication and characterization

#### a. Probe fabrication and assembly

Figure 2(A) shows a 50-μm-thick processed microfluidic probe after release from the grinding tape. The pronounced color difference between probe base and shank is due to the high density of interconnecting lines along the shank. Details of the tapered shank tip with the IrO_x_ electrodes and the fluidic outlet ports are provided in the scanning electron microscopy (SEM) micrographs in Fig. 2(B&C). Both outlet ports shown in the inset of Fig. 2(B) are connected to a single fluidic channel. The array of sealed etch openings above this fluidic channel are clearly visible in Fig. 2(C). Figure 2(D) illustrates the cross-section of the probe tip along X-Y, as indicated in Fig. 2(A), generated by focused ion beam (FIB) etching. It reveals an 11-μm-wide fluidic channel in the Si substrate buried under a 10-μm-thick stack of SiO_x_/SiN_y_, as well as interconnection lines under the topmost passivation stack, and the via between metallization and the IrO_x_ electrode. For FIB processing, the probe surface has been covered by a protective layer made of Pt and carbon.

An assembled microfluidic probe with PTFE tubing and PI ribbon cable is shown in Fig. 3(A). It clearly indicates the high flexibility of the cable and shows in the background its ZIF interface to be connected to the probe PCB (Fig. 3(B)). The two sets of electrodes arranged at pitches of 60 and 85 μm are easily distinguished in the inset of panel A. The fluidic functionality of the Si-based neural probe is proven by injecting water through the fluidic channel that forms a small droplet around the two fluidic outlets.

A completely assembled, multifunctional optrode attached to a custom-designed holder for the stereotaxic arm is shown in Fig. 3(B). A connection with a low dead volume between the microfluidic probe and the glass syringe is realized using thin PTFE tubings (ID 300 μm, OD 760 μm) and an interconnect union.

#### b. Electrical characterization

Figure 4(A) summarizes the absolute impedances |*Z*| of a sample of 16 electrodes of a 32-channel probe measured over the frequency range from 100 Hz to 40 kHz. The 30-μm-large circular IrO_x_ electrodes show on average a |*Z*| value of 168 ± 9 kΩ (n = 32) at a frequency of 1 kHz.

**Figure 4:**
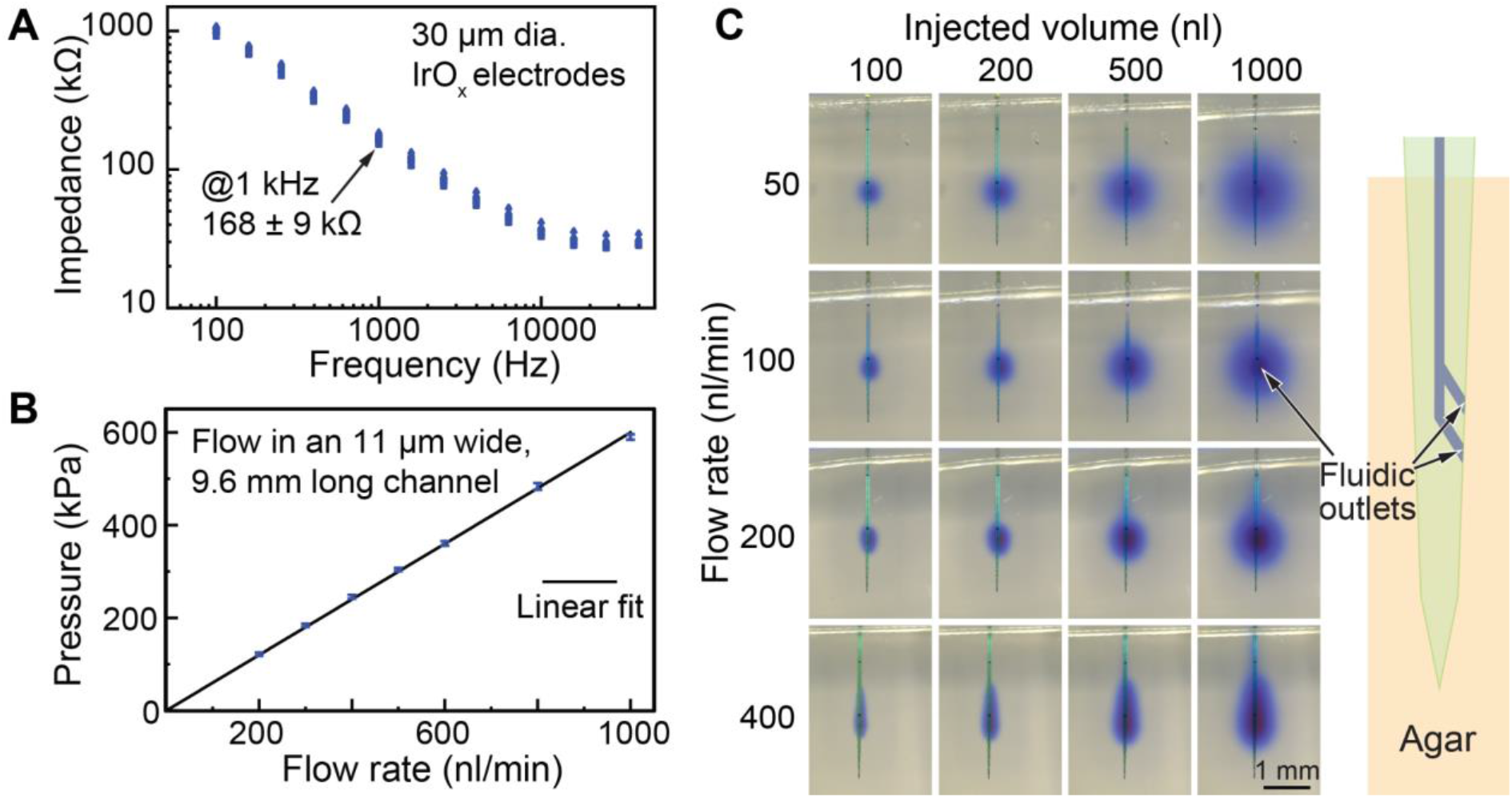
(A) Absolute impedance |Z| of a sample of 16 IrO_x_ electrodes of a 32-channel probe measured in saline. The mean value of all electrodes (n = 32) at 1 kHz is 168 ± 9 kΩ. (B) Pressure measured in the fluidic tubing as a function of applied flow rate for a probe with an 11-μm-wide and 9.6-mm-long microfluidic channel. (C) Injections of dye in agarose gel at different flow rates to observe liquid distribution. Pictures just after injecting 100, 200, 500, and 1000 nl of dye are presented for each flow rate. Fluidic outlets open to the right of the probe, as shown in the probe schematic on the right.

#### c. Fluidic characterization

Figure 4(B) shows a representative pressure measurement as a function of flow rate for a probe with an 11-μm-wide and 9.6-mm-long channel. As expected for a laminar flow inside the PTFE tubing and the probe channel, the relationship between the flow rate and measured pressure difference is linear with a flow rate of 167 nl min^−1^ at 100 kPa (= 1 bar). For applications which require a higher flow rate per applied pressure, the size of microfluidic channels can be increased during fabrication by a prolonged XeF_2_ etching time.

#### d. In vitro injections into agarose gel

Snapshots of dye distribution after injecting 100, 200, 500, and 1000 nl of dye at flow rates of 50, 100, 200, and 400 nl/min are shown in Fig. 4(C). At a low flow rate of 50 nl/min, the dye distribution around the two channel outlets is mainly isotropic and largely dominated by diffusion. In contrast, a significant backflow along the shank is observed for the highest flow rate of 400 nl/min. At the intermediate flow rates of 100 and 200 nl/min a moderate backflow resulting in a slightly elliptical dye distribution can be noticed.

### 3.2 In vivo experiments

#### a. Observation of in vivo neuromodulation via bicuculline application in an anesthetized rat

In order to demonstrate the injection via the fluidic channel and the electrophysiological recording in vivo, we locally injected a pharmacological agent known to induce a strong effect on neural activity. An acute implantation was performed, with bicuculline injected while recording neural activity with the fluidic probe. We observed a clear increase in neural activity after the injection (Fig. 5). This is in line with the well-characterized effect of bicuculline, which as a GABA receptor antagonist, causes increased neural activity and epileptic seizures (Huffman & McFadin, 1972; Pongrácz et al., 2013). The increase in activity started shortly after injection (20-70 s post-injection). This is comparable to the results of Pongrácz et al., who also found the effect to start quickly (30-60 s) post-injection (Pongrácz et al., 2013). Temporally, the increase in neural activity gradually developed from channels closest to the fluidic ports, to those farthest away. This effect could be either due to diffusion of the drug away from the port or from activated units recruiting distal neurons in synchronized seizure activity.

**Figure 5:**
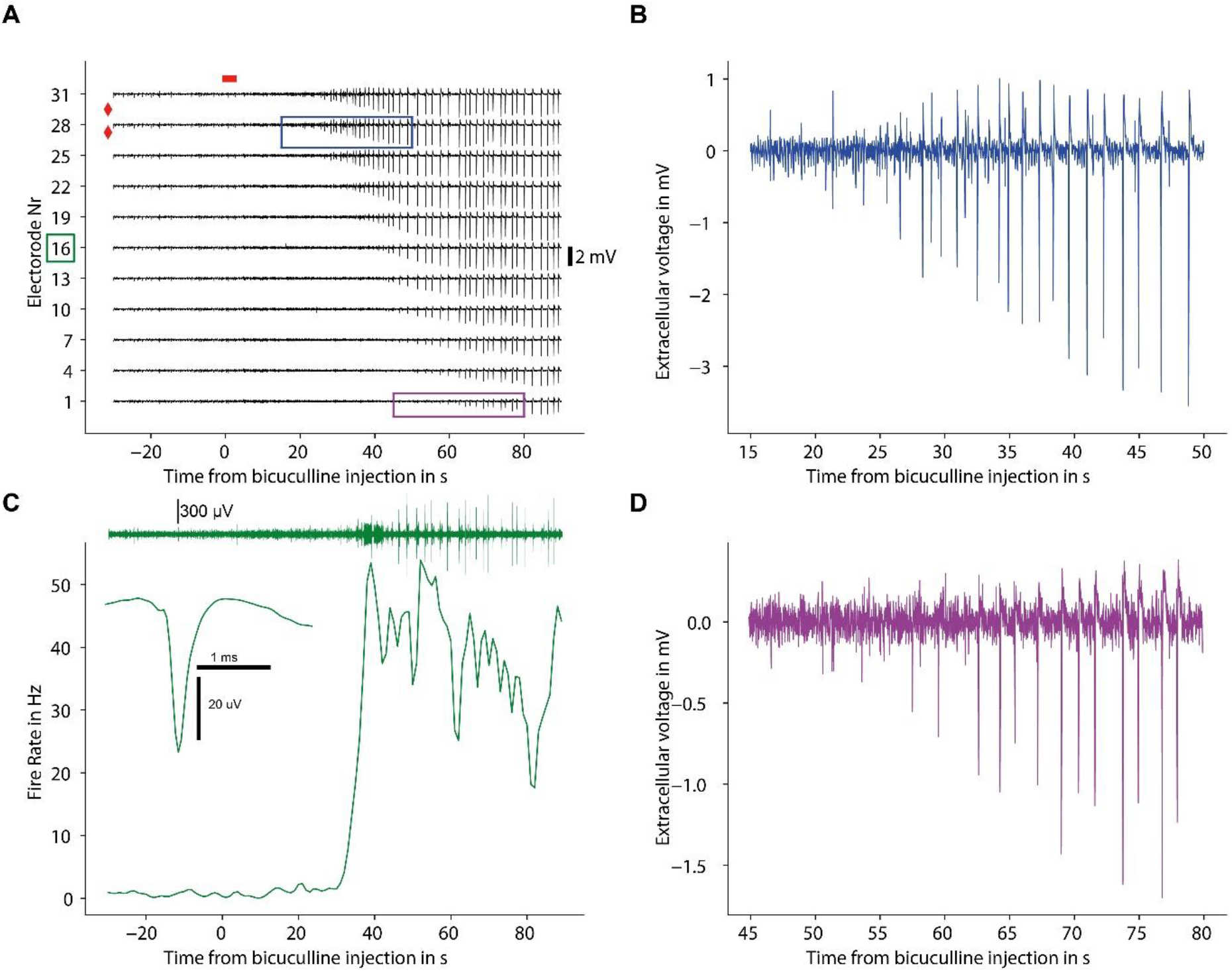
Acute implantation and bicuculline injection. (A) LFP activity measured with electrodes spanning the distance 390 μm (electrode 32) and 1.9 mm (electrode 1) from the injection site. The red diamonds represent the fluidic port location, the red rectangle represents the time span of injection. We observed epileptic activity shortly after the injection starts at all electrodes. (C) Firing rate of a neuron recorded from channel 16 before and after the bicuculline injection. The firing rate increases ~30 s post-injection. (B&D) Selected close-ups from panel A; LFP from channels 28 and 1. The increase in activity occurs earlier in channel 28 (~25 s post-injection) compared to channel 1 (~55 seconds post-injection).

#### b. Observation of in vivo neuromodulation via optogenetic activation in a freely moving rat

As ChR2 is a relevant tool for interrogating neural circuits in behavioral experiments (Tye & Deisseroth, 2012), we tested the feasibility of the multifunctional optrode for chronic implantation with viral injection of ChR2, optogenetic stimulation and electrophysiological recording. We applied PL-configured optrode, which has two injection ports and one fiber targeting PL, and recording electrodes equally distributed in both PL and IL. ChR2 was injected through the microfluidic channel of the probe during the implantation surgery. After waiting for sufficient viral expression (6 weeks), we recorded from the chronically implanted rat during blue light stimulation. We observed neural responses to blue light stimulation. As expected by stimulating ChR2 expressing neurons, there was a significant increase in the firing rate (Wilcoxon-signed rank test, Bonferroni adjusted, p < 0.001) in neural signals collected from electrodes 22-26 (Fig. 6(F)). This response was restricted to units near the upper parts of the shank (spanning 510 μm). This is expected; viral expression should be centered at the injection region near the fluidic outlets (at 1845 μm and 2045 μm from the tip). Histological analysis of the viral expression (Fig. 6(D)) also confirmed a small area of expression (ca. 600 μm dorsoventral spread), comparable to the range of ChR2-responding units found from the neural recording. A further restriction of optogenetic activation was accomplished by the fiber placement, which allowed targeted light stimulation to the upper area of the shank. We also tested different stimulation frequencies. We observed a slight increase in firing rate from responding units when stimulating with higher frequencies up to 40 Hz, which is expected since ChR2 firing rate can follow stimulation frequencies up to 40 Hz (Grossman et al., 2011; Jackman et al., 2014).

**Figure 6A-I:**
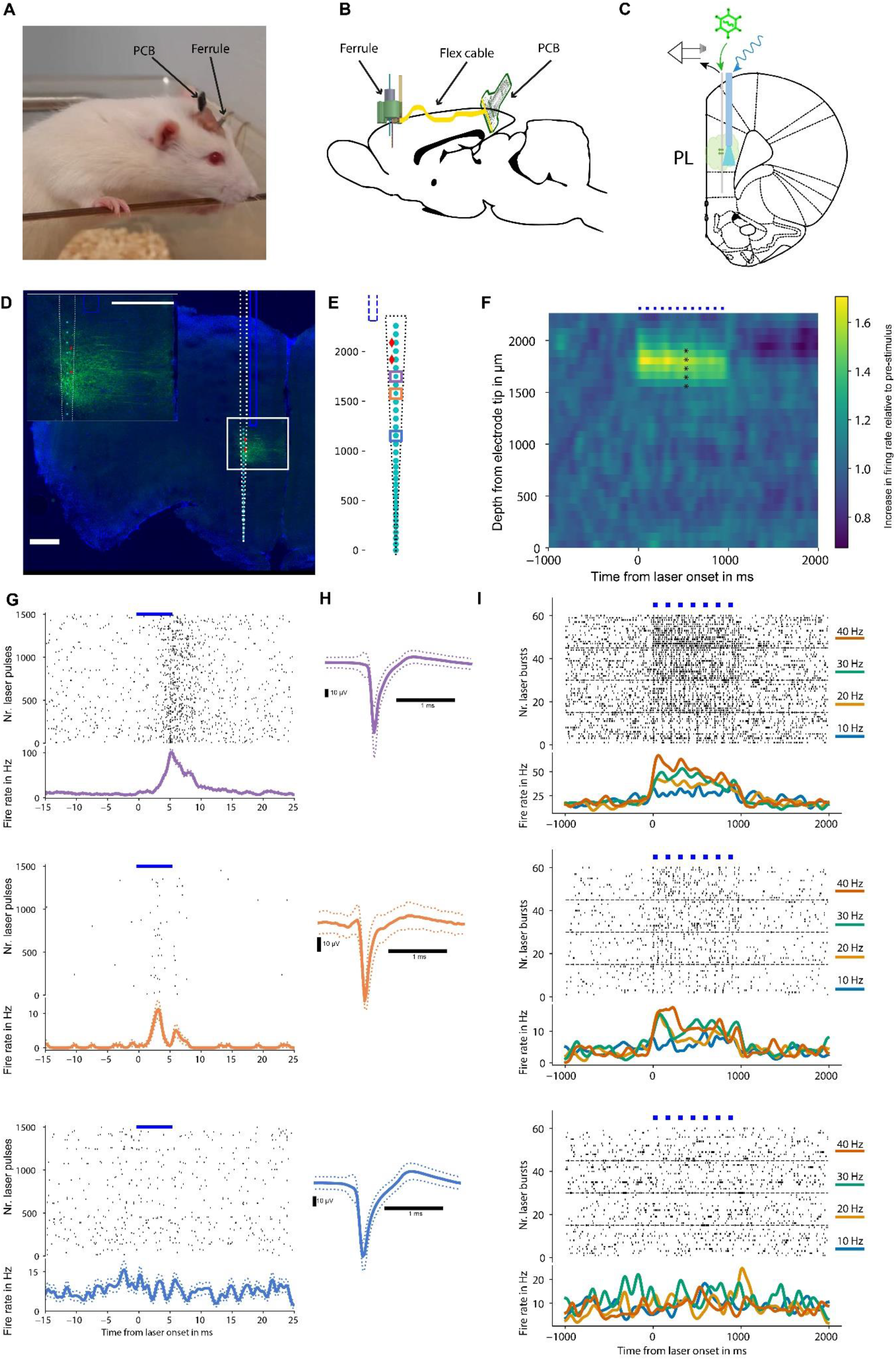
Chronic implantation and optogenetic stimulation: (A) Wild-type Sprague Dawley rat implanted with the fluidic optrode, 1 week post-implantation. The implant has comparable size to a conventional electrode or fiber implantation. The PCB is cemented at roughly a 40° angle onto the posterior part of the skull. (B) Configuration of the implant elements on the rat skull. The 3D assembly unit, the lower part of the ferrule, flex cable, and PCB board (until the ZIF connector) are embedded in dental cement. (C) Schematic of the viral injection, light activation and recording in mPFC. (D) Histology of a rat chronically implanted with a PL-configuration multifunctional optrode, which was injected via the fluidic probe with AAV5-hSyn-ChR2-eYFP. Scale bar = 500 μm. We observed a well-restricted area labelled with eYFP. The inset represents a close-up of the eYFP (representing opsin)-expressing area, with well-labelled cell bodies. (E) Schematic of the probe used for chronic implantation; the fiber is shown in blue next to the probe shank, with fluidic ports represented by red diamonds and electrodes shown in cyan. (F) Overview of neural activity with light stimulation, showing the increase in firing rate relative to 1 s pre-stimulation; * indicates electrodes with significant change of spiking activity There was an optogenetic response to blue light pulsed stimulation (1s bursts of 5 ms wide pulses represented by blue dots), which is restricted to the upper electrodes on the probe shank, located in the PL (matching the observed area of expression in Fig. 6(D)). (H&I) Spiking activity and waveforms of units recorded from channel 26, 24 and 19 (highlighted in Fig. 6(E) in respective colors), which was located in the PL of a freely moving rat. Channels 26 and 24 (purple and orange) are example units which showed an optogenetic response to 5-ms-wide blue light pulses. Such units were observed in several electrodes located in PL. The lower part of the shank (example unit recorded from channel 19 shown in blue) did not show an optogenetic response. Injection and stimulation was well restricted to the target area. (I) Comparison of the firing rate of units recorded from channel 26, 24 and 19 when stimulated with frequencies of 10, 20, 30 or 40 Hz. We observed a greater effect on firing rate for higher frequency stimulation on the optogenetically activated units (shown on channels 26 and 24).

#### c. Histology reveals spatially precise expression pattern

Histological analysis confirmed viral injection to PL. We observed eYFP expression around the target area of injection (Fig. 6(D)) spreading over an area of ca. 0.6 mm^2^ (DV spread .6 mm, ML spread 1 mm) in a 50 μm slice taken from anterior-posterior position of +3.7 mm to Bregma. Since the eYFP was injected via the viral construct that also contains ChR2, areas that express eYFP should indicate areas that also express the excitatory opsin ChR2. Notably, there was no backflow observed, as sometimes seen with traditional injection methods (Fig. S1). The small dorsoventral spread (ca. 600 μm) of opsin expressing neurons was in line with findings from the stimulation and recordings. As there was only viral expression in a restricted area of the brain, it was expected that only these opsin-expressing neurons positioned adequately near the fiber tip would respond to blue light stimulation.

## 4. Discussion and conclusions

In this paper, we presented a multifunctional optrode comprising a silicon-based microfluidic neural probe arranged close to an optical glass fiber in a miniature assembly. The microfluidic probe is equipped with an 11-μm-wide fluidic channel, and 32 IrO_x_-based recording electrodes (diameter 30 μm) on a tapered probe shank with a length, thickness, and maximum width of 7.5 mm, 50 μm, and 150 μm, respectively. Our technology enables high spatial specificity between fluidic ports and recording electrodes, with the freedom to control their size and position according to the application. Without any modification in the process, one can extend the probe design to a multi-shank array with multiple fluidic channels in the desired configuration. The use of a standard optical fiber as a waveguide facilitates the use of a wide range of optical wavelengths and power amplitudes. All elements of our multifunctional optrode are easily interfaced to external devices using commercial connectors. Optical connection to the fiber is achieved via a ferrule, electrical connection to the recording system via a TDT ZIF-clip connector on a custom-made PCB, and fluidic connection to a syringe via standard fluidic connectors.

With a modest total weight of 0.97 g, the system is suitable for chronic implantations in rodents. We demonstrated the utility of our multifunctional optrode for injecting a viral vector through the integrated microchannel, chronic implantation for several weeks to lend sufficient time for viral expression, and photostimulating neurons near the injection site while recording neural activity both at and adjacent to the injection area in an awake freely moving rat. We measured several units which responded to light stimulation, while units outside the injection area showed no response. The units which responded to an optical stimulation reflected the opsin-expressing area found in the histological analysis. The histology of this implanted rat showed a specific and restricted area of infection, which can prove advantageous to other injection methods which can result in larger viral spread post-injection. This advantage is attributed to small fluidic injection ports, and the integration of an injection modality into the optrode enabling a single insertion surgery for optogenetics. We established the multifunctional optrode as a tool for specific viral injection and optogenetic manipulation with electrophysiological recording in freely moving animals.

Fluidic channels can also be employed to apply anti-inflammatory drugs which may improve the longevity of chronic recordings, or to apply pharmacology while performing optogenetic experiments and allow for various constellations of technical applications. We demonstrated a pharmacological application with our microfluidic probe via local injection of the GABA antagonist bicuculline in an anesthetized rat, and measured its effect on neuronal activity by recording before, during and after the injection. As a proof of principle, we demonstrated an expected bicuculline effect that occurred first at units near the point of injection, and with time spread to units farther away.

Our multifunctional optrode uses standard glass fibers as a suitable light guide. However, these fibers (diameter 225 μm) are larger than our tapered microfluidic probe (maximum 150×50 μm2). Using smaller diameter fibers (Eriksson et al., 2021) or incorporating waveguides (Schwaerzle et al., 2017) on the microfluidic probe would further reduce the brain damage but might come at the cost of higher optical losses and lower output power. Integration of light sources such as LEDs (Ayub et al., 2020; Wu et al., 2015) or laser diodes directly on the microfluidic probe would further enhance the versatility of our device and would facilitate scaling the system to multiple light sources while maintaining a small implant size.

## Acknowledgements

The research leading to these results has partly received funding through the DFG-funded Priority Programme SPP 1926 (Next Generation Optogenetics) under grant number RU 869/5-1 and DI 1908/6-1, and the BrainLinks-BrainTools, Cluster of Excellence funded by the German Research Foundation (DFG, grant number EXC 1086). The authors gratefully acknowledge technical support by the RSC team at the Department of Microsystems Engineering (IMTEK) regarding cleanroom fabrication, the Laboratory for Nanotechnology at IMTEK for valuable help with SEM imaging, and the Freiburg Center for Interactive Materials and Bioinspired Technologies (FIT) for focused ion beam preparation.

## Authors contribution

Conceptualization, P.R. and I.D.; Methodology, K.S. and Z.J.; Analysis, K.S., Z.J., A.S.; Investigation, K.S., Z.J. and A.S.; Writing – Original Draft, K.S. and Z.J.; Writing – Review & Editing, P.R., O.P. and I.D.; Visualization, K.S., Z.J. and A.S.; Supervision, P.R., O.P. and I.D.; Funding Acquisition, P.R. and I.D.

