## Supplementary Figures for "Multifunctional optrode for opsin delivery, optical stimulation, and electrophysiological recordings in freely moving rats"

### Supplementary data

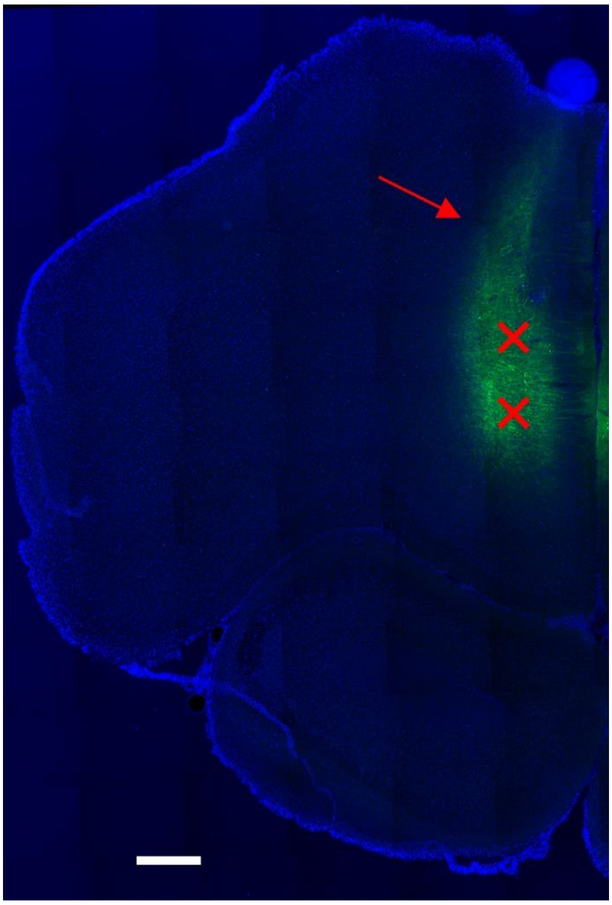

*Figure S1: Histology of a rat injected with hSyn-eYFP-ChR2 via a picospritzer and micropipette with 25-50 µm diameter tip. 500 nl of virus (250 nl at two DV depths) was injected at 100 nl/min. Injection target was the PL with stereotaxic coordinates: +3.24 mm AP/0.6 mm ML/ -2.2, -2.8 mm DV, marked with red crosses. One can observe eYFP expression extending dorsoventrally from ca. –0.5 DV to -3.4 DV. The red arrow indicates the undesired backflow of the virus extending dorsally above the injection area from ca. -0.5 DV to -1.5 DV. Scale bar: 500 µm.*

#### Multifunctional optrode holder

Custom-designed, 3D-printed holders facilitate the final assembly, testing, and storage of the multifunctional optrodes. The optrode is inserted into a holder during its assembly with the ferrule-terminated fiber and the PCB (Fig. S2(A)). The finished optrode stays in the holder, which is then stored in a plastic box for safekeeping. A small chamber can be introduced into the holder from the bottom and fixed using small magnets. In its uppermost position, the chamber encloses the fiber and the microfluidic probe shank. During storage, a chamber filled with 0.6% agar gel is installed to protect the microfluidic probe outlets from dust contamination (Fig. S2(B)). The holder retains the optrode during final testing before in vivo implantation. This involves optical power and electrical noise measurements. Optrode's ferrule is interfaced to an external laser, and optical power from the tip of the fiber is measured using a power meter (Fig. S2(C)). Additionally, the functionality of electrodes is verified by immersing the electrodes and a ground wire into saline solution, and connecting the PCB to the recording system. A low noise-level confirms a good, low-impedance electrode. Saline-filled chambers are installed into the holder for this purpose (Fig. S2(D)).

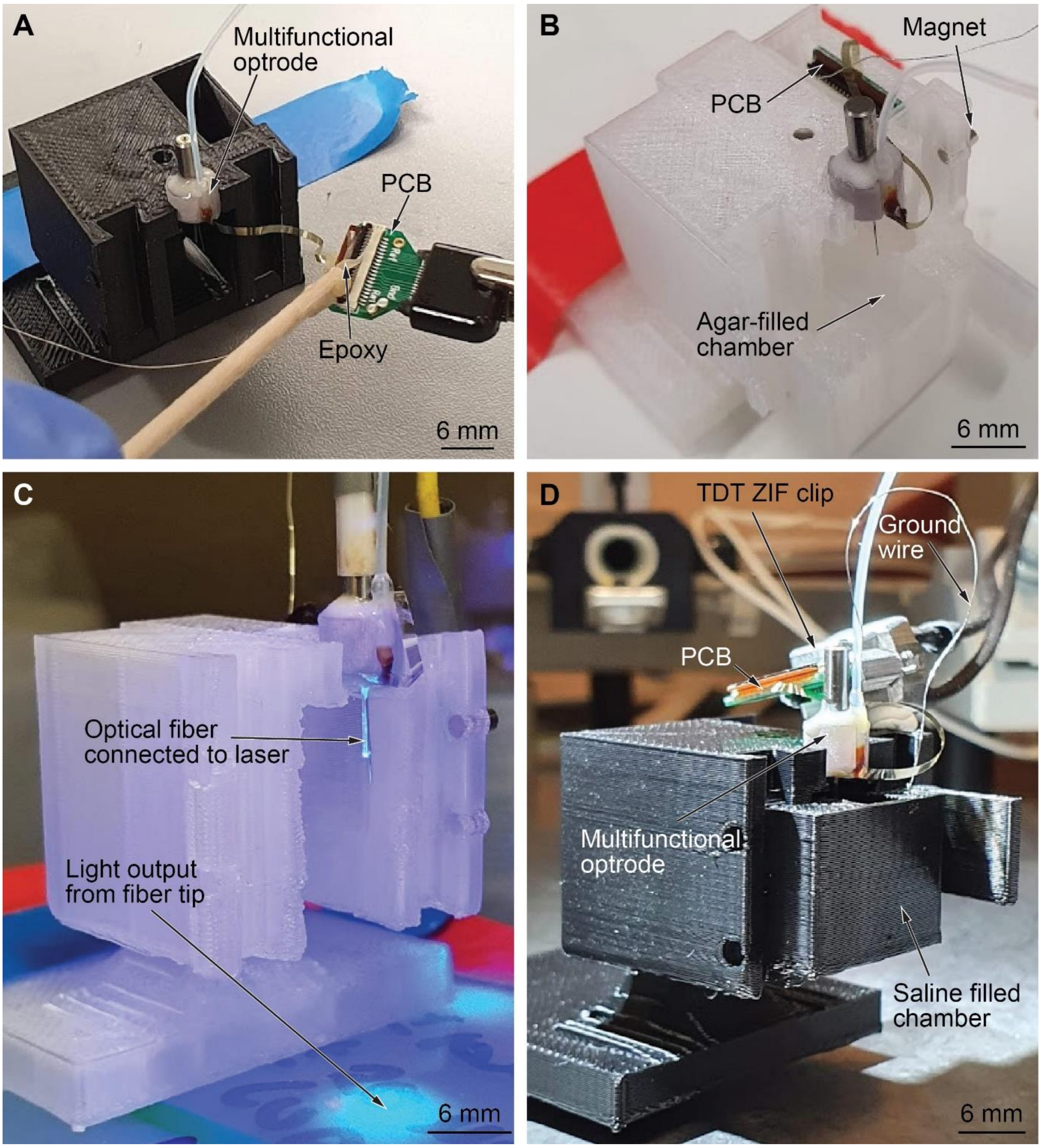


*Figure S2 Multiple applications of versatile optrode holder as (A) support during assembly of ferrule-terminated fiber and PCB into multifunctional optrode, (B) storage holder after complete assembly, and as carrier during (C) optical power testing and (D) electrode testing.*
